# Complexity of brain dynamics as a correlate of consciousness in anaesthetized monkeys

**DOI:** 10.1101/2021.08.17.456627

**Authors:** Nicolas Fuentes, Alexis García, Ramón Guevara, Roberto Orofino, Diego M. Mateos

## Abstract

The use of anaesthesia is a fundamental tool in the investigation of consciousness. Anesthesia procedures allow to investigate different states of consciousness from sedation to deep anesthesia within controlled scenarios. In this study we use information quantifiers to measure the complexity of electrocorticogram recordings in monkeys. We apply these metrics to compare different stages of general anesthesia for evaluating consciousness in several anesthesia protocols. We find that the complexity of brain activity can be used as a correlate of consciousness. For two of the anaesthetics used, propofol and medetomidine, we find that the anaesthetised state is accompanied by a reduction in the complexity of brain activity. On the other hand we observe that use of ketamine produces an increase in complexity measurements. We relate this observation with increase activity within certain brain regions associated with the ketamine used doses. Our measurements indicate that complexity of brain activity is a good indicator for a general evaluation of different levels of consciousness awareness, both in anesthetized and non anesthetizes states.

## 1 Introduction

The last few decades have witnessed significant advances in our understanding of the neural basis of consciousness, as new technological developments in brain imaging and electrophysiological methods allow for a precise spatio-temporal sampling of neural activity. [1]. Fundamental for this field is the idea that some features of brain activity correlate with different states of consciousness. The search for such features is crucial as it endows the field of consciousness with an empirical, falsifiable tool to quantify the elusive nature of subjective experience. Therefore, it is a common practice in consciousness research to investigate altered states of consciousness (e. g., coma, deep sleep stages, vegetative states or epileptic seizures), as the corresponding activity during such states largely differs from normal brain activity.

Anaesthesia is one of the most important brain states investigated in the framework of conscious studies, as it allows for a partial or total suppression of consciousness in a safe, controlled way. Indeed, unconsciousness is one of the features of general anaesthesia. It can be indirectly assessed by the lack of clinical responsiveness such as movement, autonomic activation, or by measurements performed using EEG sensors connected to the patient. Different administration protocols lead to different depths in anaesthesia that could in principle correspond to differences in brain dynamics. However there is no single measurement (except for the lack of brain electrical activity) by which anaesthesiologist can objectively asses that a patient under general anaesthesia is unconscious.

Quantifiers based on information theory have proven to be effective in distinguishing between different brain states, such as sleep stages [2–4] and in the detection of epileptic seizures [5, 6]. These quantifiers do not require a large amount of signal pre-processing (in some cases they can be even directly applied to the raw signal) making them faster to implement in real time [7]. These types of indices had also been used to characterise brain states in the domain of psychiatric and neurodegenerative diseases research [5, 6, 8]. Within the particular field of anaesthetics research, measures such as Permutation Entropy [9] have been used to quantify the effect of sevoflurane on EEG signals, obtaining better results than classical measures such as the Bispectral Index (BIS) [10–12]. Lempel-Ziv complexity has been used to study EEG signals in patients under the effects of propofol [13]. Another study compares different entropic measures in patients under anaesthesia induced by GABAergic agents [14].

In this work we used Electrocortical (ECoG) signals datasets provided by the Artificial Intelligence Laboratory of the University of Riken [15]. The database consists of recordings from four macaques under the effects of four different anaesthetics schemes: propofol, ketamine, medetomidine and medetomidine-ketamine. The recorded brain activity was acquired within a controlled anaesthetic environment, providing accurate data sets which allows us for a precise analysis of the different possible states of consciousness.

As it is known, not all anaesthetics act similarly on the physiology of the central nervous system (CNS), so it is to be expected that EoCG signals have different dynamics depending on the drug used. Therefore, claiming that a single quantifier is optimal for studying states of consciousness for all anaesthetics could be wrong. To address this problem, we propose to analyse EoCG signals with three different information measures, which focus on studying unique characteristics of the signals. The first measure was the Shannon Entropy, which is a global quantifier of the system’s uncertainty. The second quantifier is Lempel-Ziv Complexity, which measures the information redundancy within the signal. The third was Fisher’s Information, which evaluates the local information contained in the signal. Each of these quantifiers focus on specific characteristics of the signal stream. The values of each quantifier were compared in each state of consciousness, and results were analysed through Complexity-Entropy planes in order to obtain supplementary information from the system. Also, we studied the distribution of information values over the cerebral cortex, and brain local variations according to different dynamical states. Results showed that two of the three quantifiers allow for the distinction of different states of consciousness. Significant differences were found between measurements for different anaesthetics schemes. Finally, it was observed that brain dynamics measurements from recovery states were significantly different from the baseline states.

## 2 Methods

### 2.1 Data

The data used in this study were taken from the open access database *Neurotycho anesthesia and sleep task* from the Artificial Intelligence Laboratory at Riken University [15]. The database consists of electrocorticographic (EoCG) signals recorded from the left hemisphere of four monkeys. The experiments consisted of measuring the electrophysiological activity of the monkeys in five different states of consciousness i) awake open eyes (AOE), ii) awake close eyes (ACE) iii) anaesthetised (AN) iv) recovery close eyes (RCE) v) recovery open eyes (ROE). For the anaesthetised states four different anaesthetics were used, ketamine (KT), methodomidine (MD), propofol (PF) and a combination of ketamine and methodomidine (KTMD). The protocol of the experiments, the doses of the drugs used and information on data acquisition is extensively detailed on the web page [16] and in [17]. Data was acquired using 128-channel EoCG equipment with 1 KHz sampling frequency. Electrodes were placed on the left hemisphere of the monkeys, with 5 mm inter-electrode distance continuously covering the frontal, parietal, temporal and occipital lobes (Figure 1A). Two of the four monkeys have electrodes on the medial side. The original signals had a duration of 5 to 20 min for each state. For this work we took 5 min of the original recording. The first and last minutes of each state were discarded to be sure that the monkey was in the state and not in a transition. The signal was downsampled from 1 KHz to 250 Hz and preprocessed by applying a passband filter between 0.5 and 100 Hz and a notch filter at 50 Hz. Figure 1B shows an example of the signals obtained after preprocessing.

**Figure 1:**
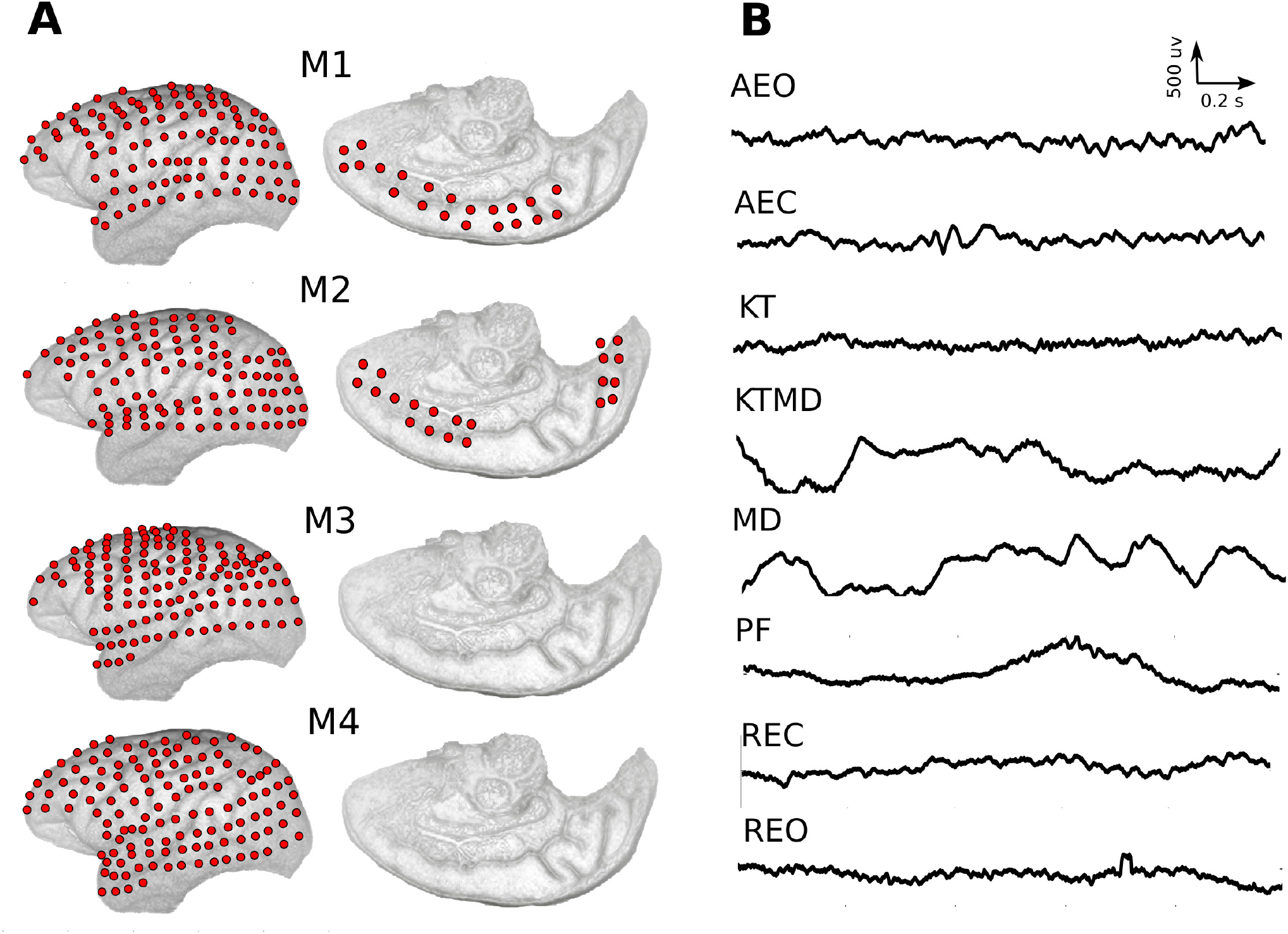
A) Electrodes distribution over the brain for the four monkeys (M1 - M4) analysed in this work. B) Example of 2 sec recording of EoCG signals, belonging to a frontal channel of monkey 1 (M1). The signals correspond to the states: awake open eyes (AOE), awake close eyes (ACE), anaesthetised under kethamine (KT), kethamine + methodomidine (KTMD), methodomidine (MD), propofol (PF), recovery close eyes (RCE) and recovery open eyes (ROE).

### 2.2 Time series discretization using ordinal pattern approach

As a pre-processing step, a discretization of the time series is performed. The study and characterisation of time series 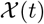 by recourse to information theory tools assume that the underlying probability distribution function (PDF) is given a priory. In the literature there are many methods to quantify continuous time series, such as binarization, histograms or wavelet, among others. However, an effective method that emerges naturally is the one introduced by Bandt and Pompe in 2002 called permutation vectors [18]. This method is based on the relative values of the neighbours belonging to the series, so it takes into account the time structure or causality of the process that generated the sequence. To understand this idea, let us consider a real-valued discrete-time series 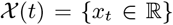, and let *D* ≥ 2 and *τ* ≥ 1 be two integers. They will be called the embedding dimension and the time delay, respectively. From the original time series, we introduce a *D*-dimensional vector 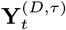:

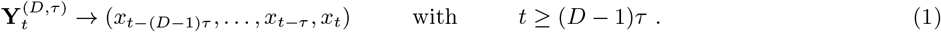

The vector 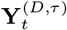 preserves the dynamical properties of the full dynamical system depending on the order conditions specified by *D* and *τ*. The components of the phase space trajectory 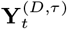 are sorted in ascending order. Then, we can define a *permutation vector*, 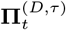, with components given by the original position of the sorted values in ascending order. Each one of these vectors represents a pattern (or motif) with *D*! possible patterns. To clarify, let us show how all this works with an example. Suppose we have a continuous series such as 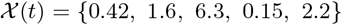 and take the parameters *D* = 3 and *τ* = 1. The embedding vectors 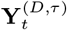 are in this case defined as 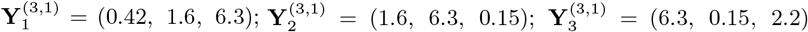, and the respective permutation vectors are 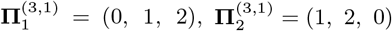 and 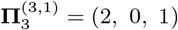.

Regarding the selection of the parameters, Bandt and Pompe [18] suggested working with 3 ≤ *D* ≤ 6 and specifically considering an embedding delay *τ* = 1. Nevertheless, other values of *τ* could provide additional information. It has been recently shown that this parameter is strongly related to the intrinsic time scales of the system under analysis [19–21].

### 2.3 Information quantifiers

In this work we used three information quantifiers to characterise brain dynamics on the basis of ECoG recordings: Permutation Shannon Entropy (PE) [18], Permutation Lempel-Ziv complexity (PLZC) [22] and Fisher Information (FI) [23]. Beyond the myriad of information measures in the literature, we particularly chose these three quantifiers due to the fact that in principle each one can extract different features of the ECoG signals. In the following sections we give a brief description of each of them.

#### Permutation Shannon entropy

Permutation Shannon entropy (PE) measures the uncertainty degree of a system. When the probability distribution of the system states is uniform (random signals) entropy tends to be maximal, while for purely deterministic signals such as periodic systems entropy is very small. Its a global measure of information, in the sense that it quantifies the information about the whole time series under consideration.

Given a time series 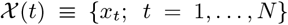, with *N* the number of observations, the Shannon’s logarithmic information measure (Shannon entropy) [24] of the associated probability distribution function (PDF), *P* ≡ {*p_i_*; *i* = 1,…, *M* } with 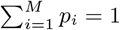, and *M* the number of possible states is defined as:

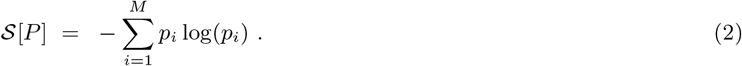

When there is total certainty that the system is in the state *i* the probability *p_i_* = 1 and this functional is equal to zero. In contrast, when the probability distribution is uniform, *P_u_* ≡ {*p_i_* = 1/*M*; ∀_*i*_ = 1,…,*M*}, knowledge about the system is minimum (all the states have the same probability) and the entropy reach its maximum.

Bandt and Pompe defined Permutation Entropy as Shannon entropy applied to the distribution of ordinal patterns [18]. Given the series of ordinal patterns 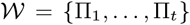, obtained from the time series 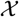 and 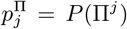, with *j* = 1,…,*D*! the probability of occurrence of the pattern Π_*j*_, the normalised permutation entropy is defined as:

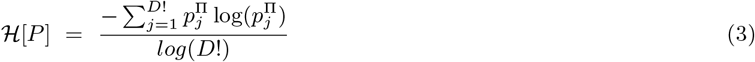

#### Lempel–Ziv complexity

To estimate the complexity of a time series 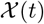 we use Lempel–Ziv complexity (LZC) [25], which is based on Kolmogorov complexity. The Kolmogorov complexity of a sequence of symbols is the minimal size of the computer program that can produce it as an output [26]. Lempel-Ziv complexity is obtained as follows. A sequence of symbols 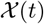 is parsed into a number 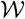 of words by considering any subsequence that has not yet been encountered as a new word. The Lempel–Ziv complexity *c_LZ_* is the minimum number of words 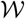 required to reconstruct the information contained in the original time series. For example, the sequence 100110111001010001011 can be parsed in 7 words: 1 · 0 · 01 · 101 · 1100 · 1010 · 001011, giving a complexity *c_LZ_* = 7. An easy way to apply the Lempel–Ziv algorithm can be found in [27]. The LZC can be normalized based in the length N of the discrete sequence and the alphabet length (*α*) as:

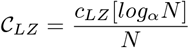

Although Lempel and Ziv developed the complexity for binary sequences, it can be used for any finite alphabet. Based on this, Zozor et al. [22] applied LZC on signals quantified by ordinal patterns, this method has the name of *Permutation Lempel–Ziv complexity* (PLZC).

#### Fisher Information

Shannon entropy 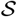 is a global information measure. To take into account local changes, it is customary use the Fisher information 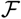 [23, 28] define as:

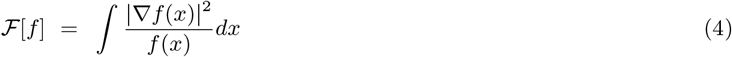

Because of the presence of a gradient operator in this equation, Fisher information is sensitive to local (differential) changes. It constitutes a measure of the curvature of the distribution *f* (*x*), so is sensitive to small, localised perturbations.

In order to compute the Fisher information of discrete time series, we adhere here to the proposal of Dehesa and coworkers [29] and define it for the discrete probability distribution *P* ≡ {*p_i_; i* = 1,…, *M*} as:

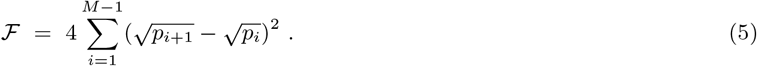

A system that has a very few possible outcomes has a very narrow PDF. The Shannon entropy of this probability distribution is close to zero (minimal uncertainty) whereas its Fisher information is maximal (very large curvature of the PDF). On the other hand, when the possible outcomes of the system have equal probability the PDF is a very flat curve. In this case, the Shannon entropy of the PDF is large whereas its Fisher information is close to zero. In other words, Fisher information and the Shannon entropy are inversely related [30].

## 3 Results

### 3.1 Comparison of different states of consciousness

We compared the information/complexity values obtained for each state of consciousness. We did this for each of the anaesthetic schemes applied (Figure 2). The bar groups represent the anaesthesia applied –ketamine (KT), medetomidine (MD), propofol (PF), ketamine-medetomidine (KTMD)– and each individual bar is the state value –awake open eyes (AOE), awake close eyes (ACE), anaesthetised (AN), recovery close eyes (RCE), recovery open eyes (ROE). Mean values and errors were calculated over all channels of all monkeys. A statistical study was performed using a linear mixed model (ANOVA). The same letters in the figures represent states with no significant differences.

**Figure 2:**
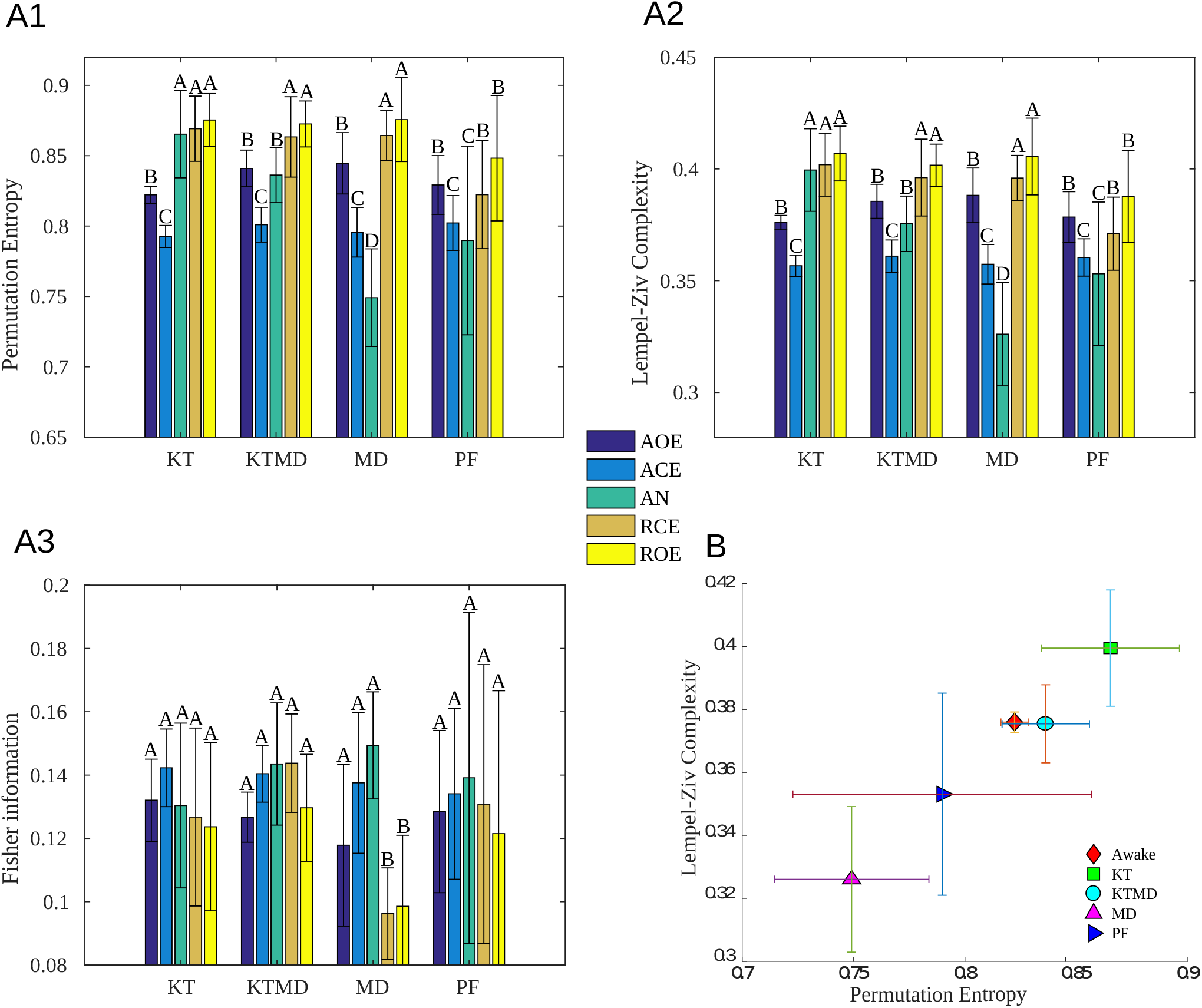
Complexity/information measures for the different states of consciousness: awake open and close eyes (AOE, ACE), anaesthetised (AN), recovery open and close eyes (RCE, ROE). The anaesthetics used were: ketanime (KT), ketanime - medetomidine (KTMD), medetomidine (MD) and propofol (PF). A1) represent Permutation entropy analysis, A2), Lempel-Ziv complexity and A3) Fisher information. The bars and errors represent the mean value and standard deviation over all channels and all monkeys. The same letters correspond to states with no significant differences. B) Analysis of ECoG signals using Complexity-Entropy plane

Figure 2 shows that permutation entropy can differentiate between awake open eyes, awake closed eyes and anaesthetised states but cannot differentiate between the anaesthetised and recovery states. The recovery states have higher entropy values than baseline states. Entropy values are lower with closed eyes in awake monkeys. Recovery entropy values are significantly higher than awake values. The values of Lempel–Ziv complexity and permutation entropy are similar. Fisher information does not distinguish differences between states. For ketamine-medetomidine, permutation entropy can differentiate between all the states. Anaesthetised values are higher than awake, but lower than recovery. Medetodo-midine shows a marked difference compared to the previous ketamine-medetomidine: entropy values are much lower in the anaesthetised than in awake and recovery; recovery values are higher than the other states. Finally, entropy values for Propofol cannot distinguish between awake closed eyes and anaesthetised states, but can discriminate between the two types of recovery states. Anaesthetised values are lower than awake and recovery. Fisher information is unable to distinguish between any states.

Generally, permutation entropy and Lempel–Ziv complexity are the only measures that can distinguish between awake open and closed eyes awake states. For all types of anaesthesia the recovery values overcomes the baseline values. None of the quantifiers can differentiate between closedand open eyes recovery states. An important point is that the permutation entropy and Lempel–Ziv complexity values are significantly different for the four types of anaesthesia. Fisher’s information values cannot differentiate between any of the states, except for MD between awake and recovering when using medetomidine.

A different way to analyse these results is through the use of complexity-entropy planes. They allow access to relevant information that is not possible to reach through the separate study of these quantifiers. These planes can be used to obtain information that is not possible by analysing the signals separately. It also allows a better visualisation of the results. In the literature there are different types of complexity-entropy planes[31–33], in this work we focus on the Permutation Lempel-Ziv complexity vs. permutation entropy plane (*LZ* × *PE*) [34]. This plane has been used to distinguish between chaotic and random signals [34], to analyse electrophysiological signals in altered states of consciousness [35] and to characterise sleep states[36].

Figure 2B shows the values of the EoCG belonging to the awake (open eyes) and anaesthetised (KT, KTMD, MD and PF) states in the Lempel–Ziv complexity–permutation entropy plane. The results show that ketamine has higher complexity and entropy values than in the awake condition. On the other hand, there is a marked decrease in entropy and complexity values for the case of propofol and medetomidine. Ketamine-medetomidine does not show significant changes with respect to the awake state. The entropy and complexity values correspond to anaesthetised states moving in areas that are above or below the awake state depending on the anaesthesia. This finding shows that each anaesthesia produces different changes in brain dynamics.

### 3.2 Differences between brain dynamics before and after anaesthetic

In Figure 3 we see that for two of the three quantifiers the recovery and baseline values are different. Because of this, a more in-depth study was carried out. We calculated the difference between the recovery and awake values. This was done for both eyes closed and eyes open (ROE-AOE and RCE-ACE). In all cases, a statistical study was performed using a paired *t*-test. Figure 3A shows the difference between the complexity/entropy values belonging to recovery and awake open eyes over all the anaesthetics applied. All anaesthetics show significant differences for permutation entropy and Lempel– Ziv complexity, but not for Fisher information. Permutation entropy and Lempel–Ziv complexity values are higher for the recovery case compared to the baseline. Ketamine presents the highest and propofol the smallest differences between states. Figure 3B shows the difference in complexity/entropy values between the awake and recovery colsed eyes states for each anaesthetic. As in the case of open eyes, permutation entropy and Lempel–Ziv complexity differences between states are significant. Ketamine–Medetomidine, have similar values than ketamine and medetomidine alone, while propofol has a lower value. These results show that although the monkey’s state is reported as ‘recovered’ from the effects of anaesthesia, in fact the monkey has not reached its initial basal state. This fact shows the importance of information quantifiers for assessing states of consciousness as they can measure relevant information which cannot be acquired by visual analysis.

**Figure 3:**
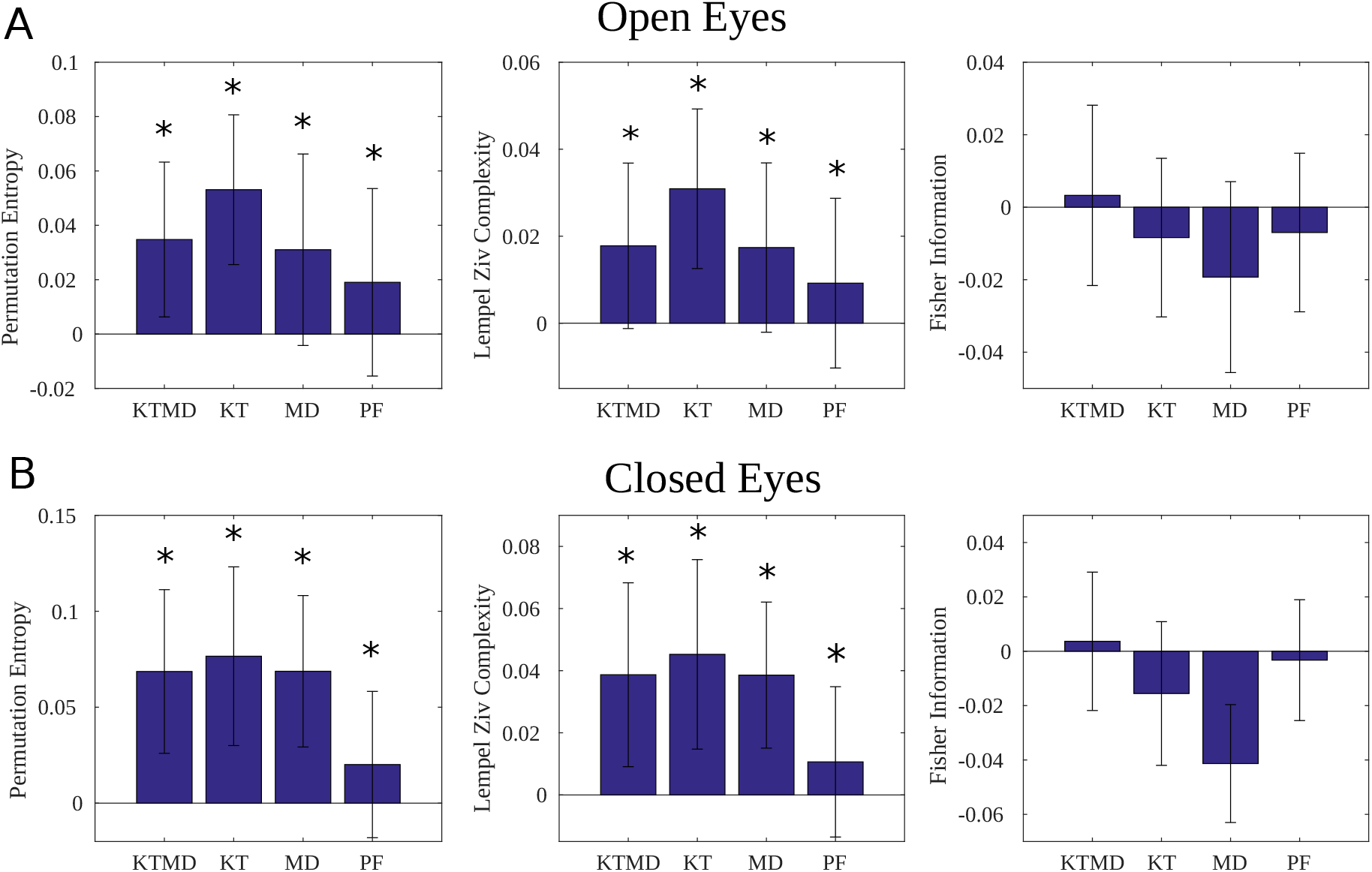
Difference between the complexity brain dynamics during recovery open eyes and awake open eyes B) Similar study for the recovery closed eyes and awake closed eyes.

### 3.3 Analysis per channel

An important aspect to study is the distribution of complexity/information values over the brain in each state of consciousness. To this end, an analysis was carried out for each channel separately. We study monkey No. 1 because it is the only one in which the four anaesthetics were applied. For this analysis the ordinal pattern parameters used were *D* = 5 and *τ* = 1.

Figure 4 shows the distribution of permutation entropy values for awake and anaesthetised states. In the awake state there is a distribution of high entropy values over the frontal-central and temporal areas, and medium values in parietal and occipital areas. Permutation entropy values decrease globally in ketamine–medetomidine, particularly in the frontal and medial areas where entropy decreased significantly. For ketamine the entropy values increased above the awake values, especially in the fronto-temporal, central and medial areas. Medetomidine shows higher values in the frontal and central area and lower values in the occipital area. Propofol presents a global decreased in entropy compared to awake states. For this state, all channels have low permutation entropy values, especially in the temporal, prefrontal and central areas (especially in the anterior electrodes).

**Figure 4:**
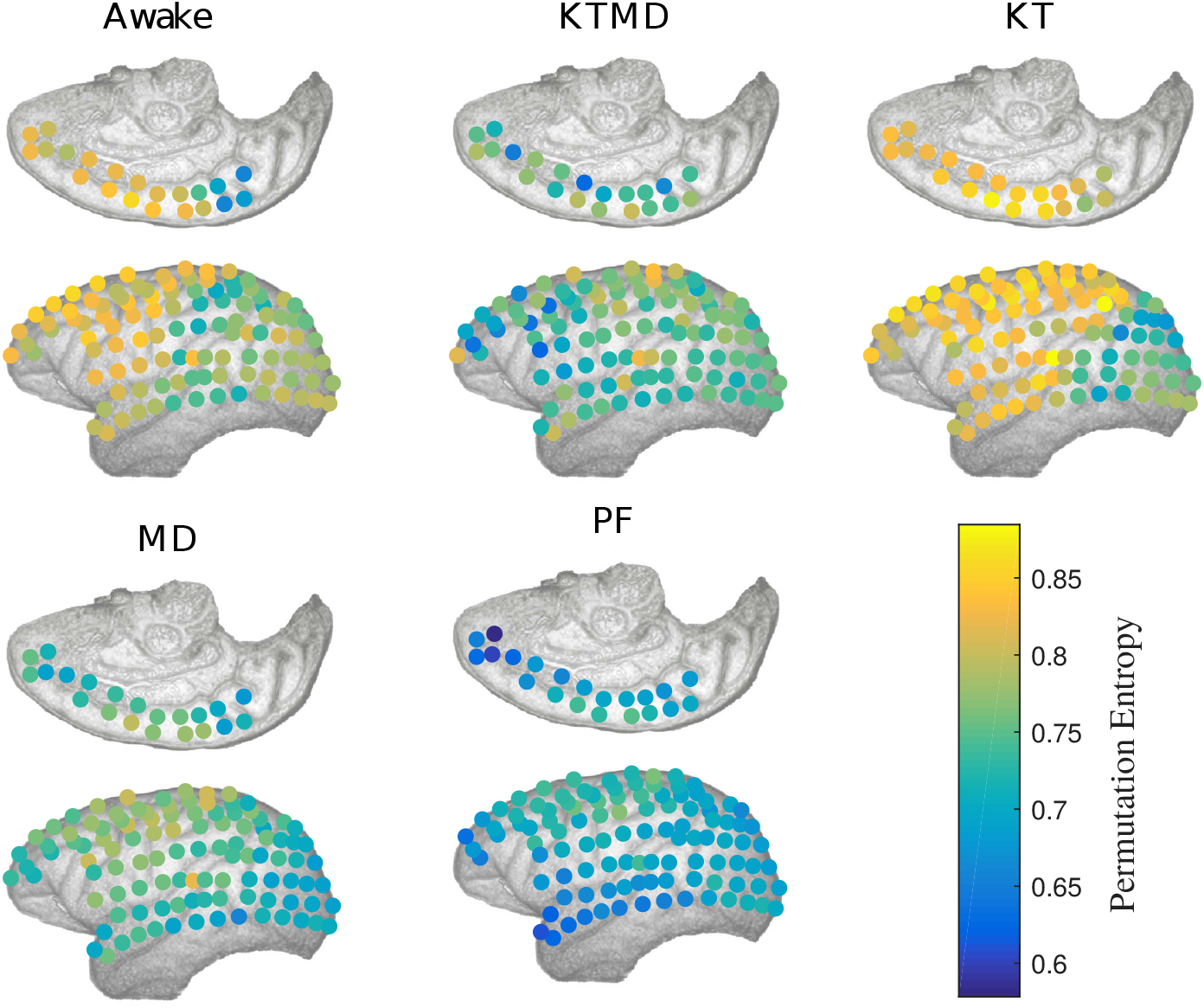
Permutation entropy analysis for each of the 125 channels belonging to Monkey 1 for the states awake (AOE) and anaesthetised (KTMD, KT, MD,PF). The ordinal pattern parameters used for the analysis were *D* = 5 and *τ* = 1.

## 4 Discussion

In this study we investigate conscious awareness in anaesthetised monkeys. We find that brain dynamics, as recorded by electrocorticography, depends on the stage of consciousness and on the type of anaesthetics used. Four types of anaesthetics schemes were used (ketamine, medetomidine, propofol and a mixture of ketamine and medetomidine) and three main stages of consciousness awareness investigated (the animal can either be awake, anaesthetised or recovering from anaesthetics; we further investigated the effect of having the eyes closed or opened).

To understand how brain dynamics correlates with the level of conscious awareness we used three measures of signal complexity: permutation entropy, Lempel–Ziv complexity and Fisher information. Taken together, these measures cover a large spectrum of signal’s features, allowing for a thorough characterisation of brain dynamics. However, even if such measures are very different, they are not independent. Indeed, we found that permutation entropy and Lempel–Ziv complexity are directly proportional so they can be used equivalently as quantifiers of the brain dynamics investigated (see Figure 2 B). This is in accordance with previous work, that shows that for the type of electrophysiological signals investigated in the current work, that is, very noisy signals, entropy and complexity tent to be equivalent [22]. This means that patterns of temporal dynamics do not add more information about the signal than what is already present in the histogram of the signal. Furthermore, it can be also inferred from our results that Fisher information measures are inversely related to both permutation entropy and Lempel–Ziv complexity (see Figure 2 A3). This is because Fisher information is a local quantifier of the signal whereas permutation entropy and Lempel–Ziv complexity are global quantifiers. As we discuss in more detail in the methods section, global and local (differential) measures are inversely related. In other words, our results suggest there are strong correlations between measures of brain activity for the three information theory metrics used. We can conclude that in the dynamic range used, electrophysiological signals can be completely characterised by any of them. So, in the following, we discuss our results in terms of permutation entropy alone, and we refer to it as “complexity”.

We found significant differences in the complexity of brain activity between different anaesthetics schemes. The general result is the following: medetomidine and propofol decrease the complexity of brain activity but ketamine doesn’t. More specifically (see Figure 2 A1): 1) Under the effect of anaesthesia, the complexity of brain activity is the lowest in the anaesthetised state than in any other state investigated, both for medetomidine and propofol. 2) This is not true when ketamine is used (alone or accompanied by medetomidine). If ketamine is used alone, the anaesthetised state has more complexity than the initial, awake state. By mixing ketamine with medetomidine, this effect is diminished.

Physiological, this can be explained in terms of the specific action of these anaesthetics. In the case of propofol, it works by binding to *GABA_A_* receptors, triggering widespread inhibition of neuronal activity. At low doses, propofol induces states of amnesia, sedation, atonia, whereas at higher doses it induces anaesthesia [37, 38]. Experimental evidence suggests propofol inhibits the ability of the brain to maintain high levels of dynamical complexity, resulting in a low-entropy state insufficient for supporting conscious awareness [39, 40]. This finding supports previous research which shows a decrease in Lempel-Ziv complexity and permutation entropy values under the effects of this anaesthetic in scalp EEG studies [41, 42].

The action of medetomidine is similar to that of propofol. This drug is a racemic mixture of two optical stereoisomers: dexmedetomidine (the active enantiomer) and levomedetomidine. It produces sympatholysis, sedation, and antinociceptive effects. It acts nonselectively on various subtypes of membrane-bound G protein-coupled *α*2-adrenoceptors. Intracellular pathways include inhibition of adenylate cyclase and modulation of calcium and potassium ion channels. This drug produces a decrease in activity of the projections of the locus coeruleus to the ventrolateral preoptic nucleus. This is an essential component of the onset of the stage of sleep non-rapid eye movement (NREM). This could be an explanation why the medetomidine values are located in the same zone of the complexity-entropy plane as those observed for subjects in the NREM state [36]. Our finding seems to be consistent with an fMRI study from subjects under different anaesthetics which showed that medetomidine had significantly lower entropy values than other anaesthetics [43].

Although sedation, analgesia and sympatholysis are produced during the administration of medetomidine, this drug must be used as a coadjuvant as it does not reach satisfactory anaesthetic conditions by itself. Association with opioids or ketamine is recommended in veterinary practice. Drug combination as adjuvants is used to reach pharmacologic effects by increasing efficacy and potency of individual drugs. There is also, as a consequence, a reduction in total doses and side effects. The use of a lower dose of both anaesthetics may be the reason why when a mixture of ketamine and medetomidine is used, the values of complexity are least variable in relation with the awake state (Figure 2 1A).

Contrary to propofol and medetomidine, brain activity under ketamine administration shows higher values of complexity in comparison with awake states. This result may be explained by the fact that ketamine acts primarily as an antagonist of glutamaterigic N-methyl-D-aspartate receptor (NMDA) [37, 44], causing widespread, light central nervous system stimulation and a state typically referred to as “dissociative anesthesia” [45, 46]. Unlike propofol, which reduces consciousness even at low doses, ketamine often produces complex conscious experiences, including hallucinations, out-of-body experiences, and dream-like, immersive experiences [44]. Ketanime’s blockade of NMDA is thought to dis-inhibit cortical neurons, causing widespread, uncoordinated excitatory activity [44, 47]. This may result in an increase in the entropy of brain activity without abolishing consciousness, artificially expanding (or at least altering) the state-space repertoire. This decorrelated signal activity, made ketamine values move towards an area where the signals with higher randomness reside in the entropy-complexity planes [32, 34]. The hypothesis in which the dynamic state of higher-than-normal entropy might correspond to a psychedelic or hallucinatory state of consciousness has become known as the Entropic Brain Hypothesis [48, 49]. Moreover, these results are consistent with other studies showing that Lempel-Ziv complexity and entropy increase under effect of ketamine [13, 47, 50].

It is important to highlight that although different anaesthetic drugs have similar observable clinical effects, the exact mechanisms of action and neurodynamic effects of these are still unknown [51]. Indeed, some medications tend to act on specific receptors while others do so on various types of receptors whereas receptors have different affinity for different drugs, even in the case of receptors from the same structure family. Furthermore, the action of anaesthetics is further complicated by their dependence on drug concentration. Indeed, drugs that act on different receptors usually do so in an overlapping way. At low concentrations they bind and activate only high affinity receptors. As drug concentrations are increased, a mass effect generates the binding and activation of receptors with less affinity. It has been proposed that many anaesthetic drugs can also act in a nonspecific and generalised way in the nervous system. They may do this by modifying the solubility of plasma membranes and its components and even by modifying the dynamics of Brownian movements of molecules that intervene in the exocytosis of synaptic vesicles. These effects may alter the precision of global electrochemical signaling [52, 53]. Increasing concentrations of a drug which generates networks activation at low doses may generate a global depressant effect at high concentrations [54].

Our study indicates that the only information quantifier that allows distinguishing between eyes opened or closed in awake states is permutation entropy. This is in agreement with recent work [55], in which it was possible to detect changes of states between eyes open and eyes closed in human scalp electroencephalography using permutation entropy. However, none of the quantifiers used were able to differentiate between the brain activity of monkeys with eyes opened or closed during the recovery state. Furthermore, the open eyes recovery state had larger complexity than any other state for all the anaesthetics used. We consider that this result is important for the evaluation of anaesthetics effects in the medical setting, since it shows that visual inspection of raw signals is not always an accurate tool. This observation leads us to propose that an analysis of states of consciousness based on more reliable quantifiers would be useful in clinical practice. Our hypothesis is that recovery values return to baseline values after a sufficiently long time. Unfortunately we do not have extensive recording time to be able to corroborate this fact. Moreover, if such records were available, it would be interesting to study the average time it takes for each anaesthetic scheme to finish its neural dynamic effect completely. However, it must be taken into account that once drug administration has been interrupted, their uncoupling from receptors and sites of action is usually variable according to their physicochemical characteristics that determine their distribution and elimination mechanisms. This implies that the effects in different places can disappear in a differential way [56–58]. Furthermore, the activation or inhibition of specific receptors at different sites can generate both a reduction or an increase in the activity of underlying networks. For example, a drug that inhibits an inhibitory network can result in increased activity in a specific region of the nervous system.

The analysis per channel showed that the variation of complexity/entropy values under the effects of anaesthetics occurs over the entire brain surface. However, there are areas which are more sensitive to changes depending on the drugs used. The general trend is that ketamine induces an increase in complexity in frontal and fronto-parietal areas whereas propofol and medetomidine produce a whole-brain decrease in complexity. This is consistent with the idea that different drugs exert their effects on diverse sites of action distributed in a heterogeneous way among the nervous system’s structures [59, 60].

Fisher information was not an efficient quantifier for distinguishing states of consciousness. This may be because changes in the ECoG dynamics are global –changes occur throughout the signal and not in localised segments, because Fisher information is a local quantifier, it is not able to detect the variations at a general level. However, these changes are detectable using global quantifiers such as permutation entropy or Lempel–Ziv complexity.

Among the limitations of this work we can mention the lack of knowledge about the subjective experiences of monkeys under the effects of ketamine. As explained above, the perception of reality seems to be altered and depends on the doses applied. We extrapolate results from reports described in humans [44, 46], but we recognise that there are limitations when comparing them with these types of animals. Another limitation is the limited number of monkeys (four) and the fact that they all received different anaesthetic regimens. Given the unavailability of such data sets, a small sample size was unavoidable. However, we hope to generalise these results in future work, in order to provide more powerful statistics.

Finally, a few words on the importance of the techniques used in this work for medical applications. The search for simple and direct quantifiers of brain activity characterising different depths of anaesthesia is fundamental for consciousness research but also crucial and in the medical setting. In particular, they serve as a guide to surgeons to evaluate the effectiveness of anaesthetics during surgical interventions. Currently, there are different indexes of anaesthesia depth commercially available (such as Fourier Transform [61], Bicoherence [10], Evoke potential [62], Burst Suppression Ratio [63] or Approximate Entropy [64]) that can be computed from electroencephalografic (EEG) recordings. The main drawback of these measures is that they often require great computational power and expensive equipment (consider, for example, the EEG anaesthesia monitor M-entropy [65], the BIS VISTA™ Monitoring System [66] or Narcotrend EEG monitor [67], among others). For this reason, the search for new types of brain quantifiers of conscious awareness, such as those proposed by us, based on faster algorithms driven by open source development, is crucial in the field of anaesthesiology. Furthermore, we showed that electroencephalography data allow for a discrimination in brain dynamics under different anaesthetics drugs regimes, so it can be useful for assessing depth of anaesthesia and for decision making scenarios in the medical setting. Measuring the complexity of brain activity have potential applications in anaesthesiology such as patient safety, quality improvement and performance analysis. Benefits could arise from a pharmacoeconomic perspective as these quantifiers may help professionals for efficient drug delivery strategies. This could result in cost reductions. Furthermore, we believe that these techniques could feed machine learning algorithms in order to develop and improve closed loop systems for automatic anaesthetic drug delivery.

## 5 Conclusion

We have found that the complexity of brain activity correlates with conscious awareness. This is in agreement with other accounts that associate consciousness with brain dynamics, and in particular to the information content or complexity of neural activity and networks. Our results show that low complexity in brain dynamics is associated with the action of anaesthetics such as propofol and medetomidine, that inhibit neural activity. We also find that this is not the case for ketamine, for which the anaesthetic increase the complexity of brain activity. This results supports the idea that the anaesthetic action of ketamine is different than other drugs, as this drug increases, rather than decrease, neural activation, and is associated to hallucination and other altered states of consciousness.

